# Genes encoding teleost orthologs of human haplo-insufficient and monoallelic genes remain in duplicate more frequently than the whole genome

**DOI:** 10.1101/2021.01.12.426466

**Authors:** Floriane Picolo, Anna Grandchamp, Benoît Piégu, Reiner A. Veitia, Philippe Monget

**Affiliations:** PRC, UMR85, INRAE, CNRS, IFCE, Université de Tours, F−37380 Nouzilly, France; Université de Paris, F-75006, Paris, France; Université de Paris, CNRS, Institut Jacques Monod, F−75006, Paris, France; Université Paris-Saclay, Institut de Biologie F. Jacob, Commissariat à l’Energie Atomique, Fontenay aux Roses, France

**Keywords:** Haploinsufficient genes, monoallelic genes, phylogeny, whole-genome-duplication, duplicat, singleton

## Abstract

Gene dosage is important is an important issue both in cell and evolutionary biology. Most genes are present in two copies in eukaryotic cells. The first outstanding exception is monoallelic gene expression (MA) that concerns genes localized on the X chromosome or in regions undergoing parental imprinting in eutherians, and many other genes scattered throughout the genome. The second exception concerns haploinsufficiency (HI), responsible for the fact that a single functional copy of a gene in a diploid organism is insufficient to ensure a normal biological function. One of the most important mechanisms ensuring functional innovation during evolution is Whole genome duplication (WGD). In addition to the two WGDs that have occurred in vertebrate genomes, the teleost genomes underwent an additional WGD, after their divergence from tetrapod. In the present work, we have studied on 57 teleost species whether the orthologs of human MA or HI genes remain more frequently in duplicates or returned more frequently in singleton than the rest of the genome. Our results show that the teleost orthologs of HI human genes remained more frequently in duplicate than the rest of the genome in all the teleost species studied. No signal was observed for the orthologs of genes localized on the human X chromosome or subjected to parental imprinting. Surprisingly, the teleost orthologs of the other human MA genes remained in duplicate more frequently than the rest of the genome for most teleost species. These results suggest that the teleost orthologs of MA and HI human genes also undergo selective pressures either related to absolute protein amounts and/or of dosage balance issues. However, these constraints seem to be different for MA genes in teleost in comparison with human genomes.

## Introduction

Gene dosage effects are an important phenomenon in cell biology that has evolutionary consequences. Indeed, in eukaryote cells, most genes are present in two copies that are transcribed and produce functional proteins. However, there are exceptions. The first outstanding exception is the case of monoallelic gene expression (MA). This is so for the majority of genes that are present on the X chromosome of eutherian mammals, genes that present a parental imprinting in eutherians, and genes encoding immunoglobulins and olfactory receptors (Chess et al., 2016). Monoallelic expression of genes is under an epigenetic control that is not well understood. For these genes, dysregulation of the mechanism(s) underlying monoallelic expression can lead to expression of both alleles, and to overexpression of the corresponding protein, and thus to severe pathologies (Horsthemke 2010). The second exception concerns haploinsufficiency. Haploinsufficiency is a biological phenomenon responsible for the fact that a single functional copy of a gene in a diploid organism is insufficient to ensure a normal biological function. Haploinsuffi-ciency is detected more frequently in essential genes than in nonessential genes in yeast (Ohnuki &Ohya, 2018). Two non-mutually exclusive theories have been proposed to explain the cause of haploinsufficiency: the “insufficient amounts” hypothesis and the gene dosage balance hypothesis (GDBH). The “insufficient amounts” hypothesis states that haploinsufficiency is the consequence of a reduced protein amount due to the loss of function of one allele, this amount being insufficient to ensure its biological function (Deutschbauer et al., 2005). This hypothesis does not explain why haploinsufficiency persisted over evolutionary time. The GDBH suggests that the phenotype caused by changes of protein level in a biological process is due to stoichiometric imbalances of protein complexes involved in cellular functions (Veitia, 2002; Papp et al., 2003). This hypothesis predicts that haplo-insufficient genes confer a biological defect when the amount of proteins is halved (such as A in a complex A-B-A) but also in excess in particular cases (such as B in the same complex (Veitia, 2002). In contrast to the “insufficient amounts” hypothesis, this hypothesis proposes an elegant explanation of the conservation of haploinsufficiency during evolution.

One of the most important mechanism ensuring functional innovation during evolution is gene duplication or the duplication of entire genome (Ohno et al., 1964; Hideki &Kondrashov, 2010). Whole genome duplication (WGD) events have been observed in all taxonomic groups: bacteria (Kuroda et al, 2001), unicellular eukaryotes (Manolis et al., 2004) and in plants (Adams et al, 2005). In vertebrates, there have been two rounds of duplication of the ancestral deuterostome genome (Mable et al., 2011). One of the striking features that characterize the teleost genomes is that they underwent an additional WGD, also called the teleost-specific genome duplication (TGD), after divergence from tetrapods (Glasauer &Neuhauss, 2014). This specific WGD event provided important additional genetic material, which strongly contributed to the radiation of teleost fishes (Ravi et al., 2008). Teleosts constitute a monophyletic group of ray finned fishes, and is the widest and most diverse group of vertebrates (Robinson-Rechavi et al., 2001;Taylor et al., 2003; Taylor &Raes, 2014; Christoffels et al., 2004). The high diversity of fish species combined with a recent complete duplication makes Clupeocephala a group of great interest for the study of complete genome duplication in the animal kingdom.

Unlike single-gene duplication events, a WGD provides all at once a large number of new genetic material, promoting an increased inter- and intra-specific diversity (Van de Peer et al, 2009, 2017). Interestingly, after WGD, all genes do not remain in duplicate with the same probability. Most models predict a rapid return of part of the duplicates to a singleton state (Maere et al, 2005), the extra-copies being rapidly pseudogenized (Sankoff et al, 2010). In particular for the rainbow trout, whose genome has duplicated one more time than that of the teleost about 100 my ago, it is estimated that about 48% of the genome remaind in duplicate, when the remaining 52% of the genome quickly returned to a singleton state (Berthelot et al, 2014).

Understanding the rules explaining why certain genes remain in duplicate when others return to singleton is a challenging issue. It has been shown that certain families of genes are more likely to remain as duplicates in all taxonomic groups studied. This is the case for transcription factors, protein kinases, enzymes and transporters (Conant et al., 2008). Recently, we showed that this is also the case for genes encoding membrane receptors and their ligands (Grandchamp et al., 2019). The first explanation that has been put forward to explain the fact that genes are more often kept in duplicate is that these molecules are involved in key functions common to all organisms. Their quantitative increase would favor these key functions because of an increase in the number of molecules produced (selection for an absolute dosage increase), and/or because of a compensation of a potential loss of function mutation of one of both copies. Another explanation is based on the respect of gene dosage balance. This is particularly so for proteins whose function is heavily dependent on interactions with partners.

In the present work, we have studied on 57 teleost species whether the orthologs of human genes known to present a monoallelic (MA) expression or to be haplo-insufficient (HI) in human remain more frequently in duplicates or returned more frequently in singleton than the whole genome in fish species or not.

## Results and Discussion

There is a mean number of 13882 human genes on 22836 (60.8%) that possess at least one ortholog in at least one teleost genome. Among them, an average of 9854 (range from 3530 to 10868) have returned in singleton, an average of 3135 (range from 2323 to 7066) remained in duplicate, and an average of 893 (range from 337 to 4772) that are in triplicate or more copies.

Concerning the 312 human HI genes, 299 (95.8%) possessed at least one ortholog in at least one teleost genome. Among them, an average of 172 (range from 47 to 199 depending on the studied species) have returned in singleton, an average of 85 (range from 68 to 122 depending on the species) remained in duplicate, and an average of 19 (range from 3 to 140) that are in triplicate or more copies. A total of 285 genes remained in duplicate (or more) in at least one species among the 57 teleost species studied. In comparison with the whole genome, this higher percentage of genes returned to singleton and remained in duplicate or more is significantly different for 55 species out of 57 (Chi Square analysis, p-value range from 0.058 to 4.2E-6), and for the 57 species studied (according to a hypergeometric test, p-value range from 0.034 to 8.5E-6). Moreover, in comparison with the whole genome as well, the higher percentage of genes that are in triplicate or more copies is significantly higher in the genomes of Rainbow trout, Brown trout, Atlantic salmon, Huchen, and Common Carp (p-value range from 1.3E-8 to 8.1E-4) but not in the genome of the other teleosts. These results suggest that the teleost orthologs of HI human genes are also subjected to selective pressures either related to absolute protein amounts and/or of dosage balance issues. This suggests that HI genes in humans undergo similar constraints in in teleosts.

Among the 285 genes that remained in duplicate in at least one teleost species, 76 genes remained in duplicate or more in at least 80% (45) of species. These genes encode more (from 3 to 38 more times) transcription factors than the whole genome: bHLH transcription factor binding (GO:0043425); RNA polymerase II activating transcription factor binding (GO:0001102); activating transcription factor binding (GO:0033613); transcription factor binding (GO:0008134); DNA-binding transcription factor binding (GO:0140297); DNA-binding transcription factor activity, RNA polymerase II-specific (GO:0000981); DNA-binding transcription factor activity (GO:0003700). This enrichment of GO is completely in accordance with the GO of HI genes previously reported (Veitia, 2002). There was no particularly representative GO among the genes in triplicate in the genome of teleost species. These results are compatible with both direct insufficiency of a transcription factor as well as with balance issues (as they are often multi-subunited complexes). Threshold effects can also be at play because of the strongly nonlinear relationships (sigmoidal or S-shaped) produced by the cooperative binding of a transcription factor to a cis regulatory sequence and the transcriptional response. Thus, depending on the concentration of transcription factor a halved dosage may not be sufficient to cross the threshold required for a normal transcriptional response (Veitia, 2002).

Concerning the 206 X human chromosome genes, 176 (82,6%) possessed at least one ortholog in at least one teleost genome. Among them, an average of 116 (range from 32 to 132 depending on the studied species) have returned in singleton, an average of 35 (range from 23 to 79 depending on the species) remained in duplicate, and an average of 7 (range from 0 to 54) that are in triplicate or more copies. Concerning the 90 genetic imprinting genes, 51 (56,7%) possessed at least one ortholog in at least one teleost genome. Among them, an average of 35 (range from 12 to 41 depending on the studied species) have returned in singleton, an average of 8 (range from 3 to 23 depending on the species) remained in duplicate, and an average of 3 (range from 0 to 20) that are in triplicate or more copies. So the teleost orthologs of human genes subjected to genetic imprinting or located on X human chromosome returned to singleton or remained in duplicate (or remain present as triplicates or more copies), in the same proportions than the whole genome.

Concerning the 580 human MA genes that are not localized on the X chromosome and that are not subjected to parental imprinting, 469 (80,9%) had at least one ortholog in at least one teleost genome. Among them, an average of 265 (range from 87 to 296) have returned to singleton, an average of 118 (range from 87 to 193) remained in duplicate, and an average of 26 (range from 4 to 160) that were found in triplicate or more copies. A total of 437 genes remained in duplicate in at least one species among the 57 teleost species studied. In comparison with the whole genome, the difference of percentage of genes remained in duplicate or more is significantly higher for 47 species on 57 (Chi Square analysis, p-value range from 0.055 to 6.5E-4), and for 50 species on 57 (hypergeometric test, p-value range from 0.044 to 6.2E-4). Moreover, in comparison with the whole genome as well, the difference of percentage of genes that are in triplicate or more copies is significantly higher in the genomes of Rainbow trout, Brown trout, Atlantic salmon, Huchen, and Common Carp (p-value range from 0,056 to 5.3E-3), not in the genome of the other teleosts. We found this result surprising. Indeed, one would have hypothesized that the teleost orthologs of MA human genes returned more frequently to singleton than the whole genome. This suggests that the phenomenon that the regulation -of epigenetic mechanism- of monoallelic expression is not likely to occur for these genes in teleosts. Morever, this suggests that the constraints to express only one allele in the human does not exist for these genes in teleost. Unlike the HI genes, there is no particularly representative GO among the MA genes.

## Material and methods

We studied 57 species of fish:
Amazon molly (Poecilia formosa), Atlantic herring (Clupea harengus), Atlantic salmon (Salmo salar), Ballan wrasse (Labrus bergylta), Barramundi perch (Lates calcarifer), Blue tilapia (Oreochromis aureus), Blunt-snouted clingfish (Gouania willdenowi), Brown trout (Salmo trutta), Burton’s mouthbrooder (Haplochromis burtoni), Channel bull blenny (Cottoperca gobio), Channel catfish (Ictalurus punctatus), Climbing perch (Anabas testudineus) Cod (Gadus morhua), Common carp (Cyprinus carpio common_carp_genome), Denticle herring (Denticeps clupeoides), Eastern happy (Astatotilapia calliptera), Electric eel (Electrophorus electricus), European seabass (Dicentrarchus labrax), Fugu (Takifugu rubripes), Gilthead seabream (Sparus aurata), Greater amberjack (Seriola dumerili), Guppy (Poecilia reticulata), Huchen (Hucho hucho), Indian glassy fish (Parambassis ranga), Indian medaka (Oryzias melastigma), Japanese medaka HdrR (Oryzias latipes ASM223467v1), Japanese medaka HNI (Oryzias latipes ASM223471v1), Japanese medaka HSOK (Oryzias latipes ASM223469v1), Jewelled blenny (Salarias fasciatus), Large yellow croaker (Larimichthys crocea), Live sharksucker (Echeneis naucrates), Lyretail cichlid (Neo-lamprologus brichardi), Makobe Island cichlid (Pundamilia nyererei), Mexican tetra (Astyanax mexicanus Astyanax_mexicanus-2.0), Midas cichlid (Amphilophus citrinellus), Mummichog (Fundulus heteroclitus), Nile tilapia (Oreochromis niloticus), Northern pike (Esox lucius), Orbiculate cardinalfish (Sphaeramia orbicularis), Pachon cavefish (Astyanax mexicanus Astya-nax_mexicanus-1.0.2), Pinecone soldierfish (Myripristis murdjan), Rainbow trout (Oncorhynchus mykiss), Red-bellied piranha (Pygocentrus nattereri), Sailfin molly (Poecilia latipinna), Sheepshead minnow (Cyprinodon variegatus), Shortfin molly (Poecilia mexicana), Siamese fighting fish (Betta splendens), Stickleback (Gasterosteus aculeatus), Swamp eel (Monopterus albus), Tetraodon (Tetraodon nigroviridis), Tiger tail seahorse (Hippocampus comes), Tongue sole (Cynoglossus semilaevis), Turbot (Scophthalmus maximus), Yellowtail amberjack (Seriola lalandi dorsalis), Zebra mbuna (Maylandia zebra), Zebrafish (Danio rerio), Zig-zag eel (Mastacembelus armatus).

These fish species diverged after complete TGD. The human genes were retrieved from ENSEMBL. The ortholog copy for each gene was established in every one of the 57 fish species. Then, in each species, the fate (singleton vs duplicate) of the entirety of the human gene orthologs was studied. Moreover, a total of 312 human genes known to be haploinsufficient were recovered from Clingene (https://www.ncbi.nlm.nih.gov/projects/dbvar/clingen/), 580 human genes known to be monoallelic (Nag et al., 2015), 206 X human chromosome genes was recov-ered for GeneImprint (http://www.geneimprint.com/site/genes-by-species) and 90 genetic im-printing genes (Carrel &Willard, 2005) and the fate of their fish orthologs was recovered. A list of human genes (GRCh38.p13) was generated using BioMart from Ensembl Genes 101. The set of human genes encoding a protein (protein_coding) is selected from the gene type filter. The selected attributes in the homologous category are the different species of teleostens listed in ENSEMBL. Only stable gene IDs were selected. A list of 22836 human genes encoding a protein is listed.

We got between 12,918 (Tetraodon) and 14,626 (Brown trout) orthologous genes by fish species (average: 13,882). This does not represent the entire genome of each fish, but allowed us to make strong statistics. Moreover, we compared the global evolution of the whole human genome that had orthologs in fishes with the specific evolution of human MA and HI genes in fish species. We studied whether these fish orthologs of MA and HI genes remained as a duplicate copy, or had return to singleton in the same proportion as whole human ortholog genes.

Both Chi Square test statistical analysis and hypergeometric analysis with Benjamini-Hochberg correction were used to test the hypothesis that teleost genes that are orthologs of human MA and HI genes remained more in duplicate than the whole genome. All the statistical tests conducted in our study were performed in R. Moreover, the Panther DataBase (http://www.pantherdb.org/) was used to study the gene ontology of teleost genes that are orthologs to human HI genes, and Fisher test with Benjamini-Hochberg correction was used to classify genes according to the family.

**Figure 1.**
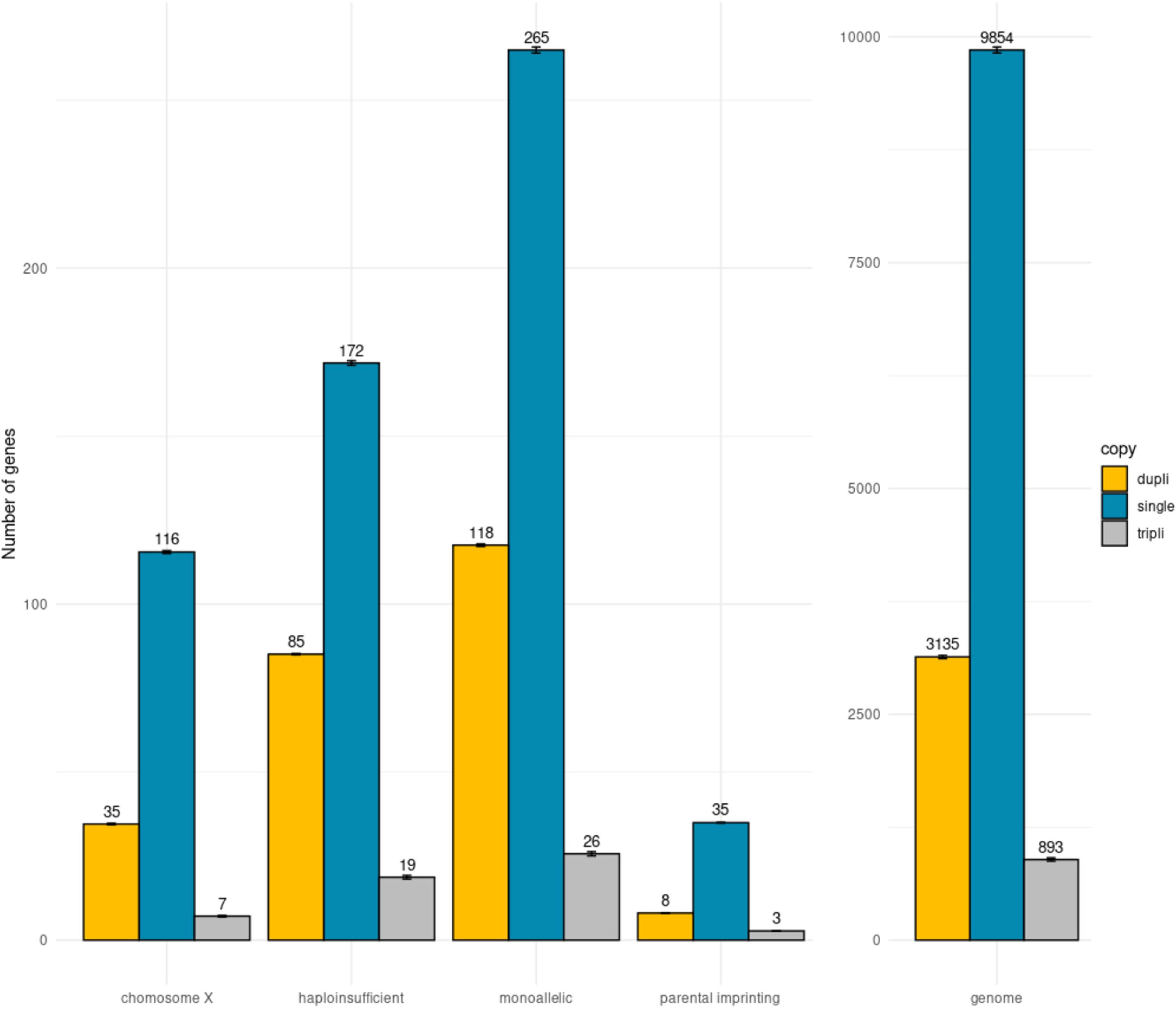
Barplot of the global distribution of the genes in each category. Teleost orthologs of human genes localized on the X chromosome, of human haplo-insufficient (HI) genes, of human genes of monoallelic expression (MA, except genes that present a parental imprinting and localized on the X chromosome), of human genes that present a parental imprinting. Right: Teleost orthologs of human genes of the whole genome. The yellow bars correspond to the genes that remained in duplicate; the blue bars correspond to the genes returned in singleton. The grey bars correspond to the genes in triplicate or more. The results are presented as mean ± SEM. * indicates a significant difference compared with the whole genome (p < 0.05).

## Acknowledgements

This work was supported by INRAE institute and by a thesis scholarship funded by the University of Tours (France). Thanks also to Alexandra Louis and Hugues Roest Crollius for helpful discussion and technical assistance.

## Suppl Data 1

Table of statistic tests for each species of teleost and for each category. Each category (HI, MA, X Chromosome, Parental imprinting) have the same construction. By column (category HI for example): (A) Species; (B) Total number of teleost genes returned in singleton. (C) Total number of teleost genes remained in duplicate; (D) Total number of teleost genes in triplicate or more copies; (E) Total number of teleost genes with a human ortholog; (F) Number of teleost orthologs of HI human genes returned in singleton; (G) Number of teleost orthologs of HI human genes remained in duplicate; (G) Number of teleost orthologs of HI hu-man genes in triplicate or more copies; (I) Total number of teleost orthologs to human HI gene; (J) Chi2 value of the repartition of HI orthologs in singleton, in duplicate or more copies in com-parison with the whole genome; (K) P-value of Chi2 test; (L) Chi2 False Discovery Rate (FDR) by Benjamini Hochberg (BH) procedure. (M) P-value of hypergeometric test between singleton and duplicate/more copies. (N) Hypergeometric FDR by BH procedure. (O) P-value of hyperge-ometric test between triplicate or more copies and in duplicate or less copies. (P) Hypergeometric FDR by BH procedure.

The same organization of columns is used for the other categories (MA, X chromosome, im-printed genes).

Concerning for HI and MA categories, the chi2 test is significant for 55/57 and 47/57 species, respectively, and the hypergeometric test is significant for 57/57 and 50/57 species, respectively, i.e. these orthologs remain more frequently in duplicate than the whole genome. For comparison between triplicate (or more copies) and duplicate (or less copies), the hypergeometric test is sig-nificant for 5/57 (salmonids and carp) and 3/57 species respectively (salmonid and carp as well).

---

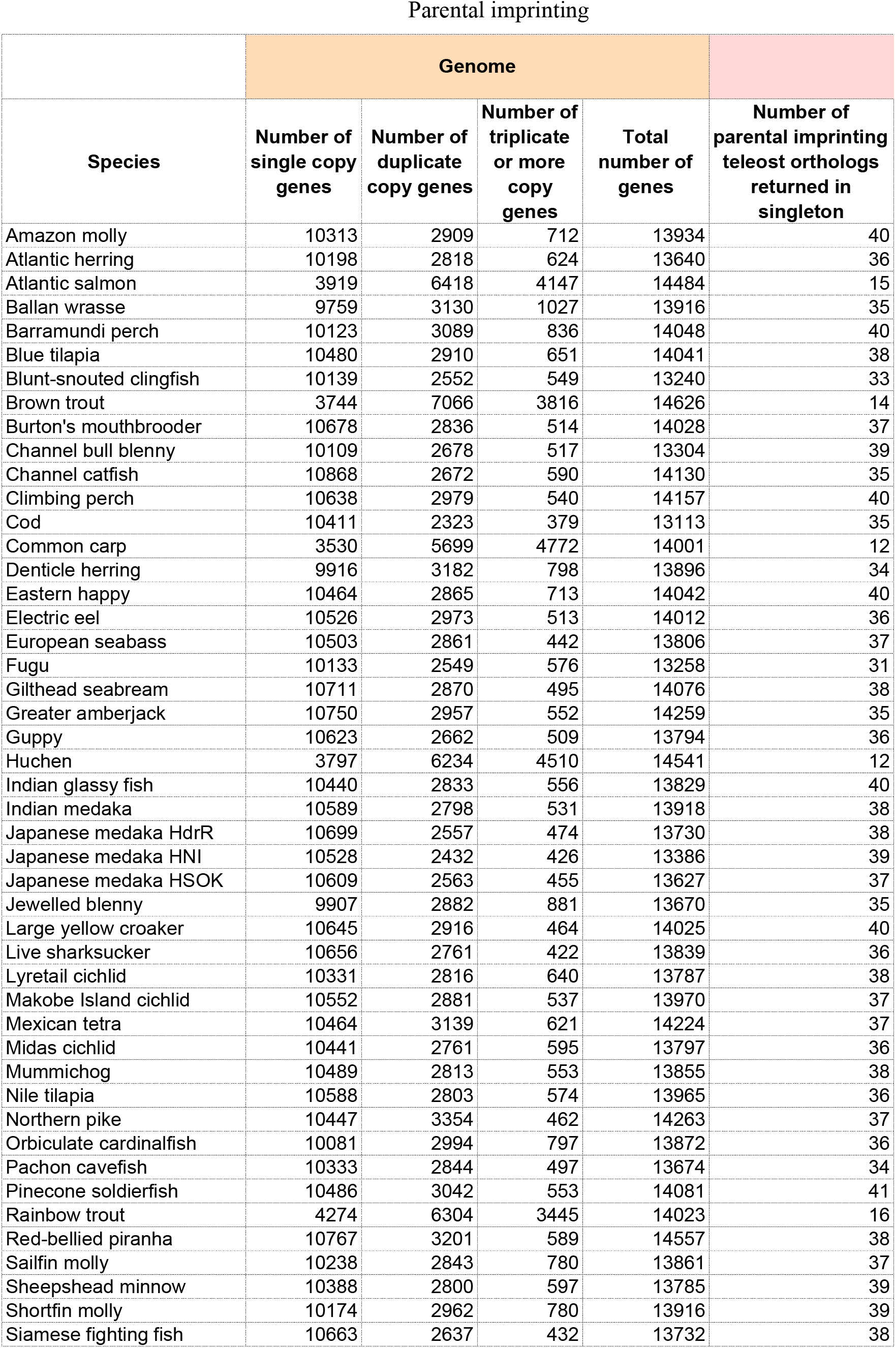
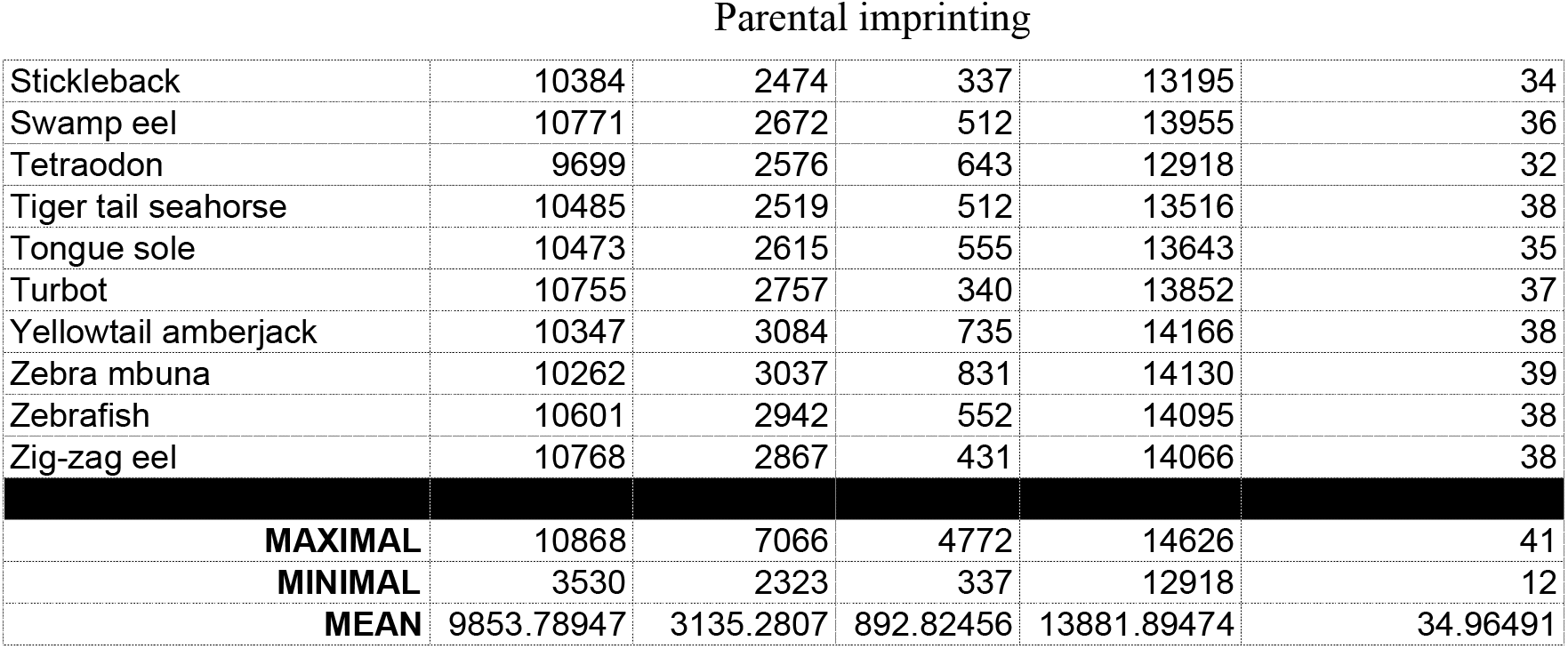

|  | Genome |  |  |  |  |
| --- | --- | --- | --- | --- | --- |
| Species | Number of single copy genes | Number of duplicate copy genes | Number of triplicate or more copy genes | Total number of genes | Number of parental imprinting teleost orthologs returned in singleton |
| Amazon molly | 10313 | 2909 | 712 | 13934 | 40 |
| Atlantic herring | 10198 | 2818 | 624 | 13640 | 36 |
| Atlantic salmon | 3919 | 6418 | 4147 | 14484 | 15 |
| Ballan wrasse | 9759 | 3130 | 1027 | 13916 | 35 |
| Barramundi perch | 10123 | 3089 | 836 | 14048 | 40 |
| Blue tilapia | 10480 | 2910 | 651 | 14041 | 38 |
| Blunt-snouted clingfish | 10139 | 2552 | 549 | 13240 | 33 |
| Brown trout | 3744 | 7066 | 3816 | 14626 | 14 |
| Burton's mouthbrooder | 10678 | 2836 | 514 | 14028 | 37 |
| Channel bull blenny | 10109 | 2678 | 517 | 13304 | 39 |
| Channel catfish | 10868 | 2672 | 590 | 14130 | 35 |
| Climbing perch | 10638 | 2979 | 540 | 14157 | 40 |
| Cod | 10411 | 2323 | 379 | 13113 | 35 |
| Common carp | 3530 | 5699 | 4772 | 14001 | 12 |
| Denticle herring | 9916 | 3182 | 798 | 13896 | 34 |
| Eastern happy | 10464 | 2865 | 713 | 14042 | 40 |
| Electric eel | 10526 | 2973 | 513 | 14012 | 36 |
| European seabass | 10503 | 2861 | 442 | 13806 | 37 |
| Fugu | 10133 | 2549 | 576 | 13258 | 31 |
| Gilthead seabream | 10711 | 2870 | 495 | 14076 | 38 |
| Greater amberjack | 10750 | 2957 | 552 | 14259 | 35 |
| Guppy | 10623 | 2662 | 509 | 13794 | 36 |
| Huchen | 3797 | 6234 | 4510 | 14541 | 12 |
| Indian glassy fish | 10440 | 2833 | 556 | 13829 | 40 |
| Indian medaka | 10589 | 2798 | 531 | 13918 | 38 |
| Japanese medaka HdrR | 10699 | 2557 | 474 | 13730 | 38 |
| Japanese medaka HNI | 10528 | 2432 | 426 | 13386 | 39 |
| Japanese medaka HSOK | 10609 | 2563 | 455 | 13627 | 37 |
| Jewelled blenny | 9907 | 2882 | 881 | 13670 | 35 |
| Large yellow croaker | 10645 | 2916 | 464 | 14025 | 40 |
| Live sharksucker | 10656 | 2761 | 422 | 13839 | 36 |
| Lyretail cichlid | 10331 | 2816 | 640 | 13787 | 38 |
| Makobe Island cichlid | 10552 | 2881 | 537 | 13970 | 37 |
| Mexican tetra | 10464 | 3139 | 621 | 14224 | 37 |
| Midas cichlid | 10441 | 2761 | 595 | 13797 | 36 |
| Mummichog | 10489 | 2813 | 553 | 13855 | 38 |
| Nile tilapia | 10588 | 2803 | 574 | 13965 | 36 |
| Northern pike | 10447 | 3354 | 462 | 14263 | 37 |
| Orbiculate cardinalfish | 10081 | 2994 | 797 | 13872 | 36 |
| Pachon cavefish | 10333 | 2844 | 497 | 13674 | 34 |
| Pinecone soldierfish | 10486 | 3042 | 553 | 14081 | 41 |
| Rainbow trout | 4274 | 6304 | 3445 | 14023 | 16 |
| Red-bellied piranha | 10767 | 3201 | 589 | 14557 | 38 |
| Sailfin molly | 10238 | 2843 | 780 | 13861 | 37 |
| Sheepshead minnow | 10388 | 2800 | 597 | 13785 | 39 |
| Shortfin molly | 10174 | 2962 | 780 | 13916 | 39 |
| Siamese fighting fish | 10663 | 2637 | 432 | 13732 | 38 |
| Parental imprinting |  |  |  |  |  |
| Stickleback | 10384 | 2474 | 337 | 13195 | 34 |
| Swamp eel | 10771 | 2672 | 512 | 13955 | 36 |
| Tetraodon | 9699 | 2576 | 643 | 12918 | 32 |
| Tiger tail seahorse | 10485 | 2519 | 512 | 13516 | 38 |
| Tongue sole | 10473 | 2615 | 555 | 13643 | 35 |
| Turbot | 10755 | 2757 | 340 | 13852 | 37 |
| Yellowtail amberjack | 10347 | 3084 | 735 | 14166 | 38 |
| Zebra mbuna | 10262 | 3037 | 831 | 14130 | 39 |
| Zebrafish | 10601 | 2942 | 552 | 14095 | 38 |
| Zig-zag eel | 10768 | 2867 | 431 | 14066 | 38 |
| **MAXIMAL** | 10868 | 7066 | 4772 | 14626 | 41 |
| **MINIMAL** | 3530 | 2323 | 337 | 12918 | 12 |
| **MEAN** | 9853.78947 | 3135.2807 | 892.82456 | 13881.89474 | 34.96491 |

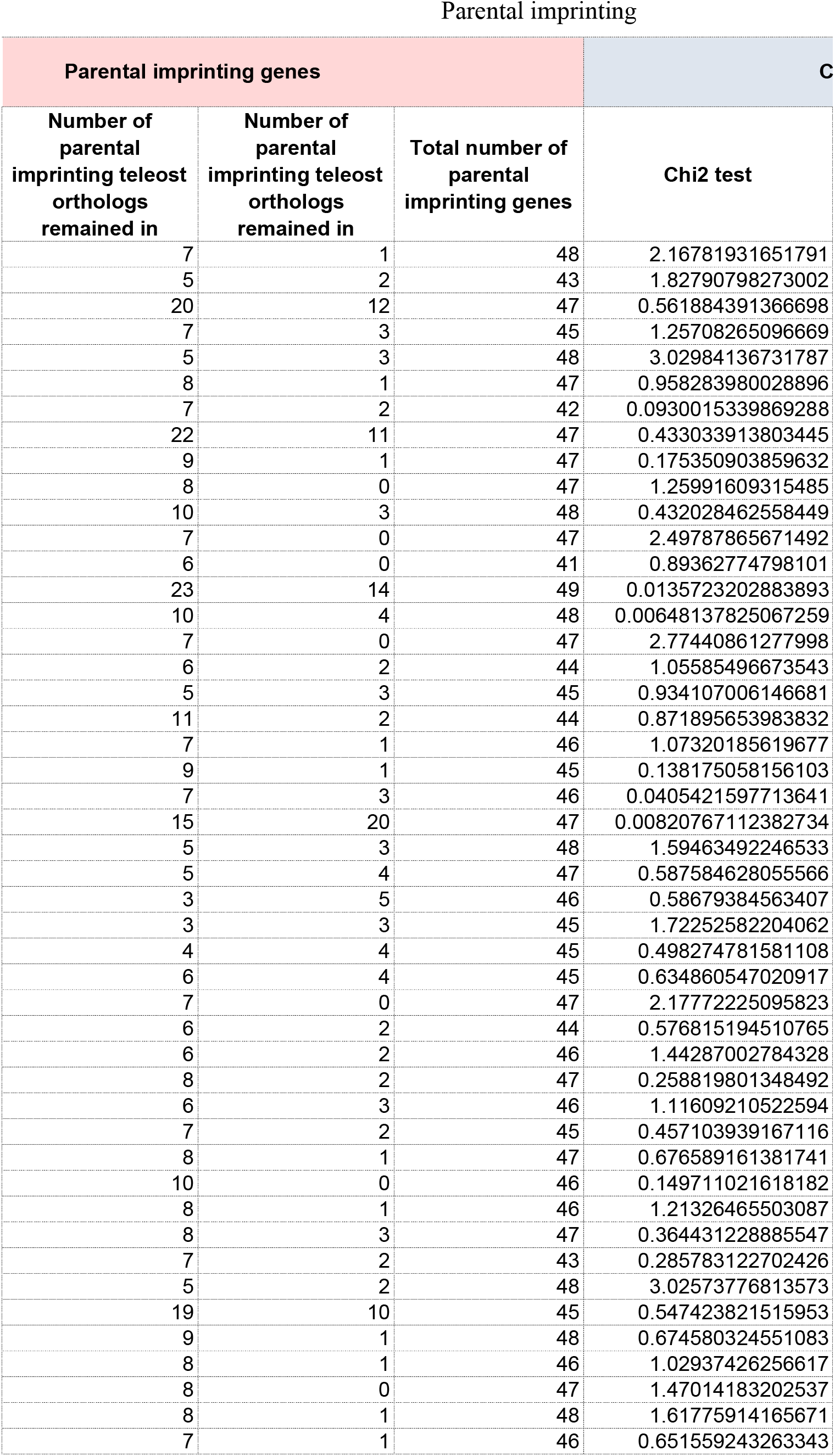
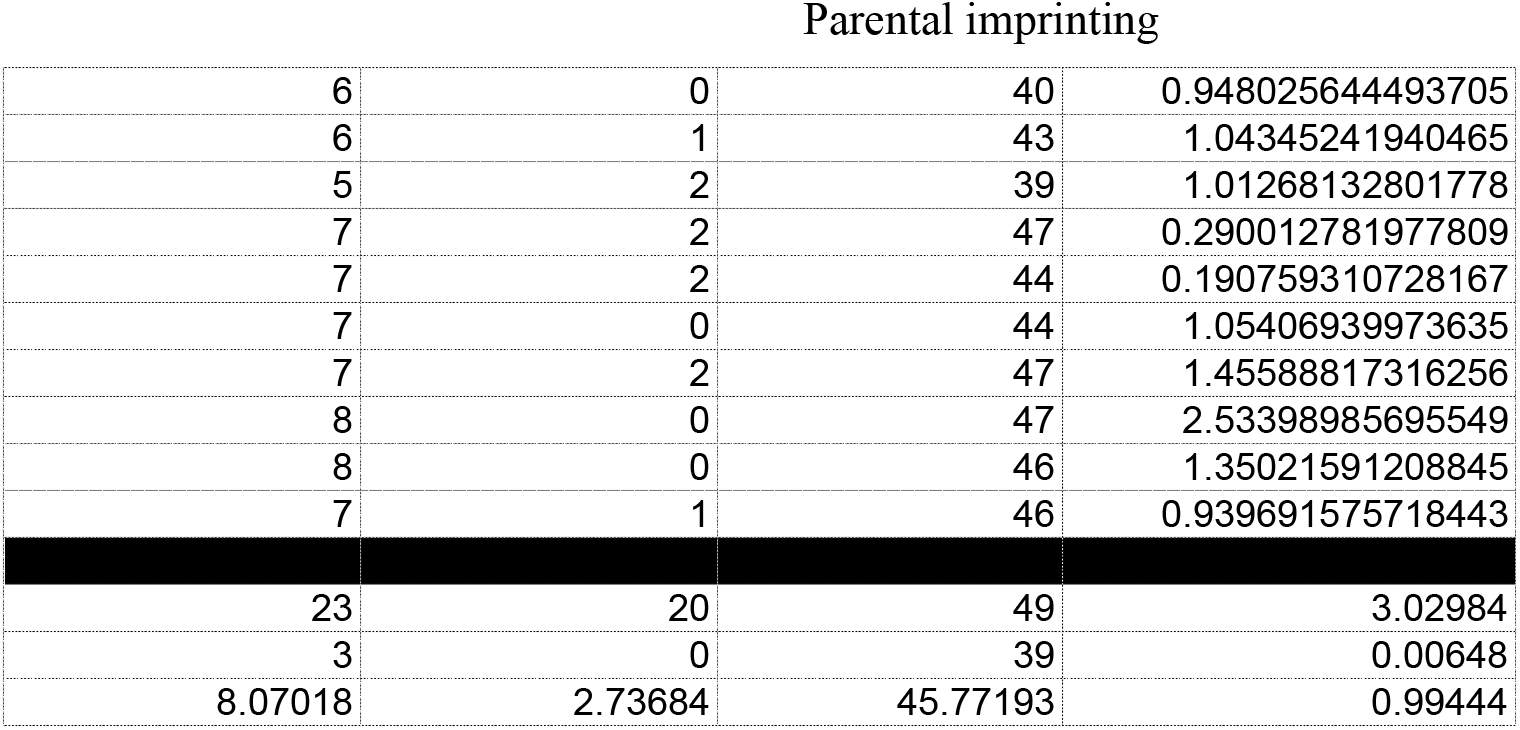

Parental imprinting
| Parental imprinting genes |  |  | C |
| --- | --- | --- | --- |
| Number of parental imprinting teleost orthologs remained in | Number of parental imprinting teleost orthologs remained in | Total number of parental imprinting genes | Chi2 test |
| 7 | 1 | 48 | 2.16781931651791 |
| 5 | 2 | 43 | 1.82790798273002 |
| 20 | 12 | 47 | 0.561884391366698 |
| 7 | 3 | 45 | 1.25708265096669 |
| 5 | 3 | 48 | 3.02984136731787 |
| 8 | 1 | 47 | 0.958283980028896 |
| 7 | 2 | 42 | 0.0930015339869288 |
| 22 | 11 | 47 | 0.433033913803445 |
| 9 | 1 | 47 | 0.175350903859632 |
| 8 | 0 | 47 | 1.25991609315485 |
| 10 | 3 | 48 | 0.432028462558449 |
| 7 | 0 | 47 | 2.49787865671492 |
| 6 | 0 | 41 | 0.89362774798101 |
| 23 | 14 | 49 | 0.0135723202883893 |
| 10 | 4 | 48 | 0.00648137825067259 |
| 7 | 0 | 47 | 2.77440861277998 |
| 6 | 2 | 44 | 1.05585496673543 |
| 5 | 3 | 45 | 0.934107006146681 |
| 11 | 2 | 44 | 0.871895653983832 |
| 7 | 1 | 46 | 1.07320185619677 |
| 9 | 1 | 45 | 0.138175058156103 |
| 7 | 3 | 46 | 0.0405421597713641 |
| 15 | 20 | 47 | 0.00820767112382734 |
| 5 | 3 | 48 | 1.59463492246533 |
| 5 | 4 | 47 | 0.587584628055566 |
| 3 | 5 | 46 | 0.58679384563407 |
| 3 | 3 | 45 | 1.72252582204062 |
| 4 | 4 | 45 | 0.498274781581108 |
| 6 | 4 | 45 | 0.634860547020917 |
| 7 | 0 | 47 | 2.17772225095823 |
| 6 | 2 | 44 | 0.576815194510765 |
| 6 | 2 | 46 | 1.44287002784328 |
| 8 | 2 | 47 | 0.258819801348492 |
| 6 | 3 | 46 | 1.11609210522594 |
| 7 | 2 | 45 | 0.457103939167116 |
| 8 | 1 | 47 | 0.676589161381741 |
| 10 | 0 | 46 | 0.149711021618182 |
| 8 | 1 | 46 | 1.21326465503087 |
| 8 | 3 | 47 | 0.364431228885547 |
| 7 | 2 | 43 | 0.285783122702426 |
| 5 | 2 | 48 | 3.02573776813573 |
| 19 | 10 | 45 | 0.547423821515953 |
| 9 | 1 | 48 | 0.674580324551083 |
| 8 | 1 | 46 | 1.02937426256617 |
| 8 | 0 | 47 | 1.47014183202537 |
| 8 | 1 | 48 | 1.61775914165671 |
| 7 | 1 | 46 | 0.651559243263343 |
| Parental imprinting |  |  |  |
| 6 | 0 | 40 | 0.948025644493705 |
| 6 | 1 | 43 | 1.04345241940465 |
| 5 | 2 | 39 | 1.01268132801778 |
| 7 | 2 | 47 | 0.290012781977809 |
| 7 | 2 | 44 | 0.190759310728167 |
| 7 | 0 | 44 | 1.05406939973635 |
| 7 | 2 | 47 | 1.45588817316256 |
| 8 | 0 | 47 | 2.53398985695549 |
| 8 | 0 | 46 | 1.35021591208845 |
| 7 | 1 | 46 | 0.939691575718443 |
| 23 | 20 | 49 | 3.02984 |
| 3 | 0 | 39 | 0.00648 |
| 8.07018 | 2.73684 | 45.77193 | 0.99444 |

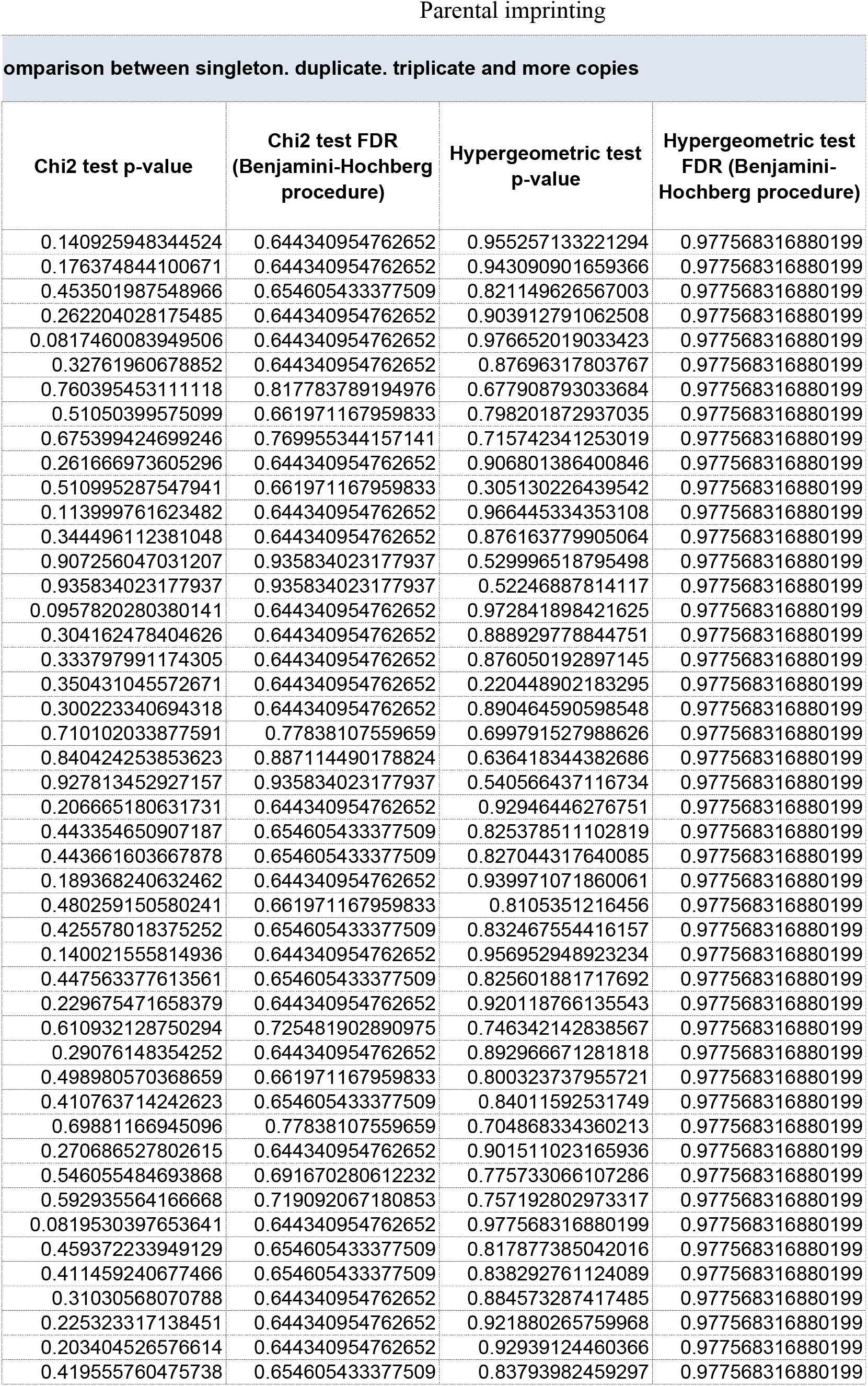
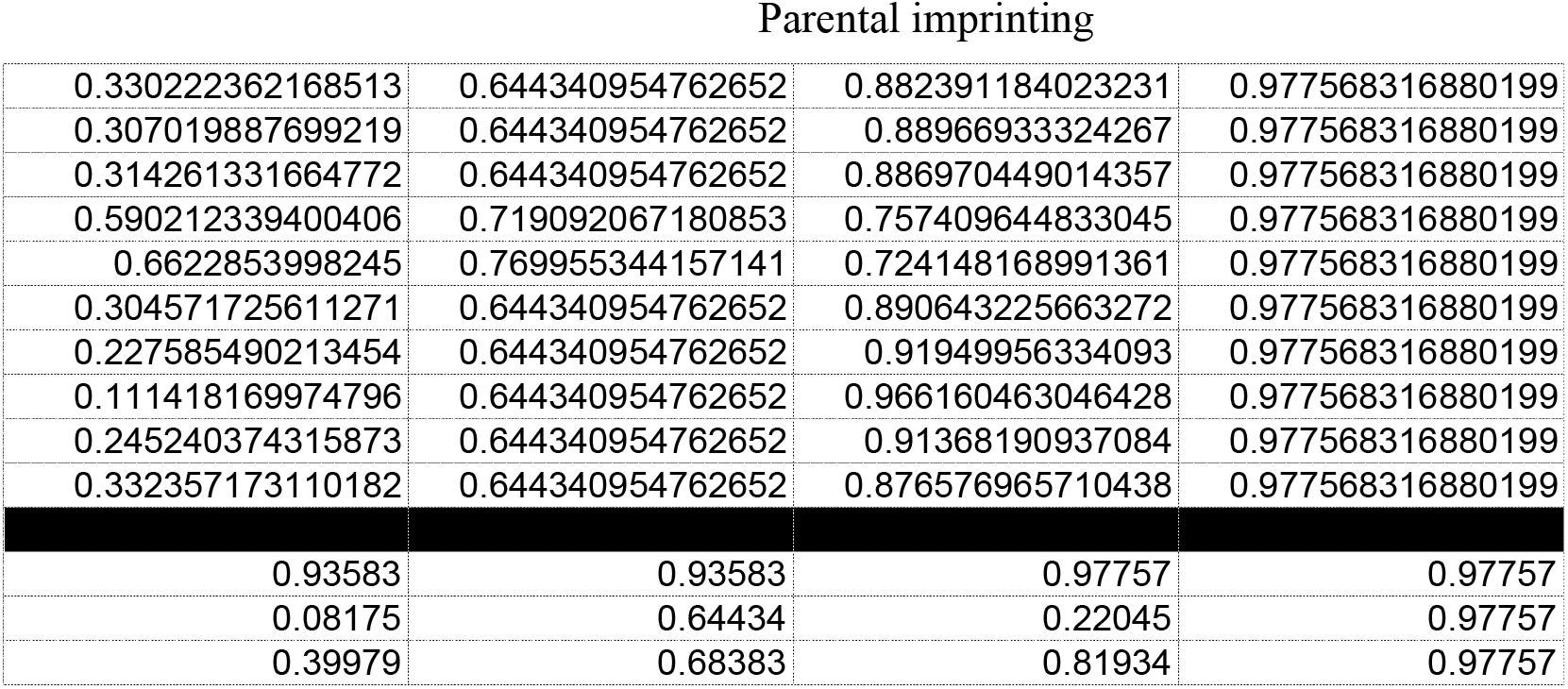

Parental imprinting
| omparison between singleton. duplicate. triplicate and more copies |  |  |  |
| --- | --- | --- | --- |
| Chi2 test p-value | Chi2 test FDR<br>(Benjamini-Hochberg<br>procedure) | Hypergeometric test<br>p-value | Hypergeometric test<br>FDR (Benjamini-Hochberg procedure) |
| 0.140925948344524 | 0.644340954762652 | 0.955257133221294 | 0.977568316880199 |
| 0.176374844100671 | 0.644340954762652 | 0.943090901659366 | 0.977568316880199 |
| 0.453501987548966 | 0.654605433377509 | 0.821149626567003 | 0.977568316880199 |
| 0.262204028175485 | 0.644340954762652 | 0.903912791062508 | 0.977568316880199 |
| 0.0817460083949506 | 0.644340954762652 | 0.976652019033423 | 0.977568316880199 |
| 0.32761960678852 | 0.644340954762652 | 0.87696317803767 | 0.977568316880199 |
| 0.760395453111118 | 0.817783789194976 | 0.677908793033684 | 0.977568316880199 |
| 0.51050399575099 | 0.661971167959833 | 0.798201872937035 | 0.977568316880199 |
| 0.675399424699246 | 0.769955344157141 | 0.715742341253019 | 0.977568316880199 |
| 0.261666973605296 | 0.644340954762652 | 0.906801386400846 | 0.977568316880199 |
| 0.510995287547941 | 0.661971167959833 | 0.305130226439542 | 0.977568316880199 |
| 0.113999761623482 | 0.644340954762652 | 0.966445334353108 | 0.977568316880199 |
| 0.344496112381048 | 0.644340954762652 | 0.876163779905064 | 0.977568316880199 |
| 0.907256047031207 | 0.935834023177937 | 0.529996518795498 | 0.977568316880199 |
| 0.935834023177937 | 0.935834023177937 | 0.52246887814117 | 0.977568316880199 |
| 0.0957820280380141 | 0.644340954762652 | 0.972841898421625 | 0.977568316880199 |
| 0.304162478404626 | 0.644340954762652 | 0.888929778844751 | 0.977568316880199 |
| 0.333797991174305 | 0.644340954762652 | 0.876050192897145 | 0.977568316880199 |
| 0.350431045572671 | 0.644340954762652 | 0.220448902183295 | 0.977568316880199 |
| 0.300223340694318 | 0.644340954762652 | 0.890464590598548 | 0.977568316880199 |
| 0.710102033877591 | 0.77838107559659 | 0.699791527988626 | 0.977568316880199 |
| 0.840424253853623 | 0.887114490178824 | 0.636418344382686 | 0.977568316880199 |
| 0.927813452927157 | 0.935834023177937 | 0.540566437116734 | 0.977568316880199 |
| 0.206665180631731 | 0.644340954762652 | 0.92946446276751 | 0.977568316880199 |
| 0.443354650907187 | 0.654605433377509 | 0.825378511102819 | 0.977568316880199 |
| 0.443661603667878 | 0.654605433377509 | 0.827044317640085 | 0.977568316880199 |
| 0.189368240632462 | 0.644340954762652 | 0.939971071860061 | 0.977568316880199 |
| 0.480259150580241 | 0.661971167959833 | 0.8105351216456 | 0.977568316880199 |
| 0.425578018375252 | 0.654605433377509 | 0.832467554416157 | 0.977568316880199 |
| 0.140021555814936 | 0.644340954762652 | 0.956952948923234 | 0.977568316880199 |
| 0.447563377613561 | 0.654605433377509 | 0.825601881717692 | 0.977568316880199 |
| 0.229675471658379 | 0.644340954762652 | 0.920118766135543 | 0.977568316880199 |
| 0.610932128750294 | 0.725481902890975 | 0.746342142838567 | 0.977568316880199 |
| 0.29076148354252 | 0.644340954762652 | 0.892966671281818 | 0.977568316880199 |
| 0.498980570368659 | 0.661971167959833 | 0.800323737955721 | 0.977568316880199 |
| 0.410763714242623 | 0.654605433377509 | 0.84011592531749 | 0.977568316880199 |
| 0.69881166945096 | 0.77838107559659 | 0.704868334360213 | 0.977568316880199 |
| 0.270686527802615 | 0.644340954762652 | 0.901511023165936 | 0.977568316880199 |
| 0.546055484693868 | 0.691670280612232 | 0.775733066107286 | 0.977568316880199 |
| 0.592935564166668 | 0.719092067180853 | 0.757192802973317 | 0.977568316880199 |
| 0.0819530397653641 | 0.644340954762652 | 0.977568316880199 | 0.977568316880199 |
| 0.459372233949129 | 0.654605433377509 | 0.817877385042016 | 0.977568316880199 |
| 0.411459240677466 | 0.654605433377509 | 0.838292761124089 | 0.977568316880199 |
| 0.31030568070788 | 0.644340954762652 | 0.884573287417485 | 0.977568316880199 |
| 0.225323317138451 | 0.644340954762652 | 0.921880265759968 | 0.977568316880199 |
| 0.203404526576614 | 0.644340954762652 | 0.92939124460366 | 0.977568316880199 |
| 0.419555760475738 | 0.654605433377509 | 0.83793982459297 | 0.977568316880199 |
| Parental imprinting |  |  |  |
| 0.330222362168513 | 0.644340954762652 | 0.882391184023231 | 0.977568316880199 |
| 0.307019887699219 | 0.644340954762652 | 0.88966933324267 | 0.977568316880199 |
| 0.314261331664772 | 0.644340954762652 | 0.886970449014357 | 0.977568316880199 |
| 0.590212339400406 | 0.719092067180853 | 0.757409644833045 | 0.977568316880199 |
| 0.6622853998245 | 0.769955344157141 | 0.724148168991361 | 0.977568316880199 |
| 0.304571725611271 | 0.644340954762652 | 0.890643225663272 | 0.977568316880199 |
| 0.227585490213454 | 0.644340954762652 | 0.91949956334093 | 0.977568316880199 |
| 0.111418169974796 | 0.644340954762652 | 0.966160463046428 | 0.977568316880199 |
| 0.245240374315873 | 0.644340954762652 | 0.91368190937084 | 0.977568316880199 |
| 0.332357173110182 | 0.644340954762652 | 0.876576965710438 | 0.977568316880199 |
| 0.93583 | 0.93583 | 0.97757 | 0.97757 |
| 0.08175 | 0.64434 | 0.22045 | 0.97757 |
| 0.39979 | 0.68383 | 0.81934 | 0.97757 |

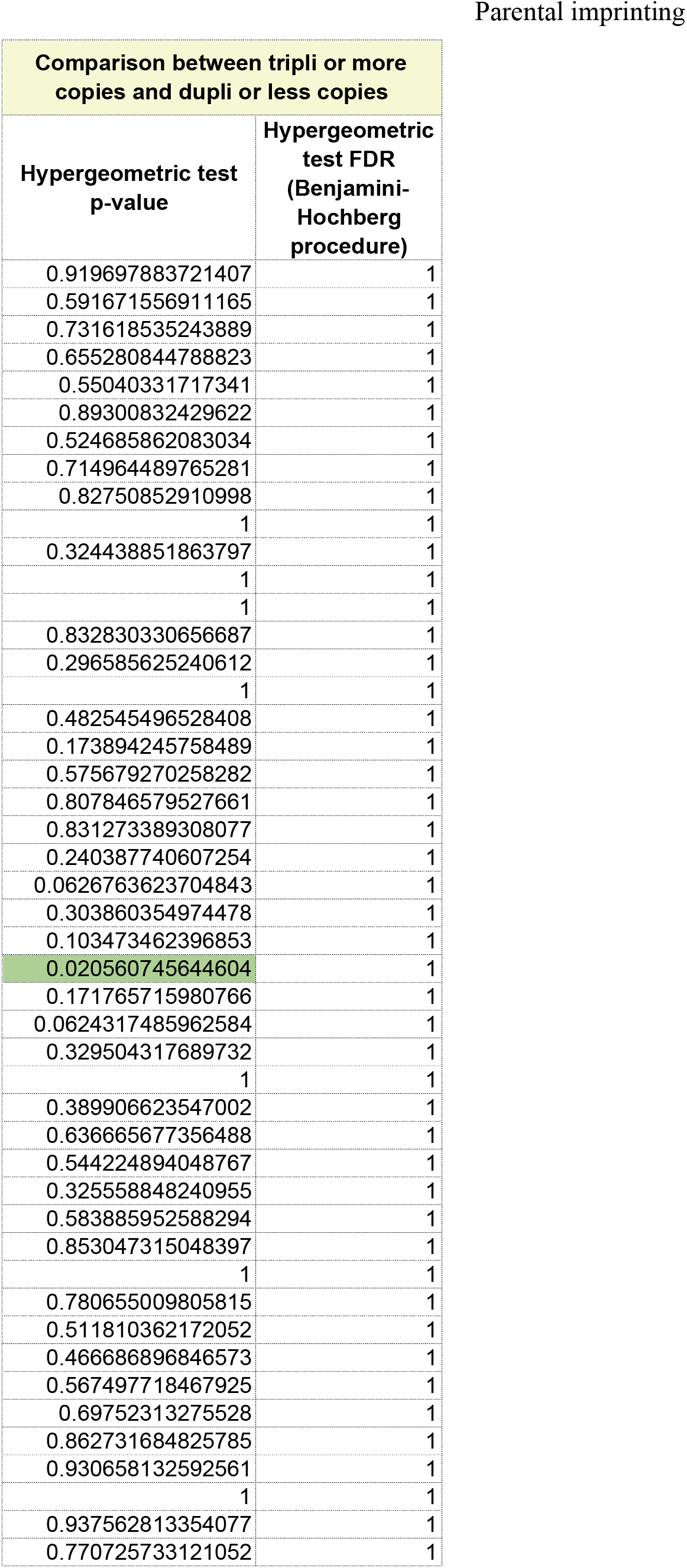
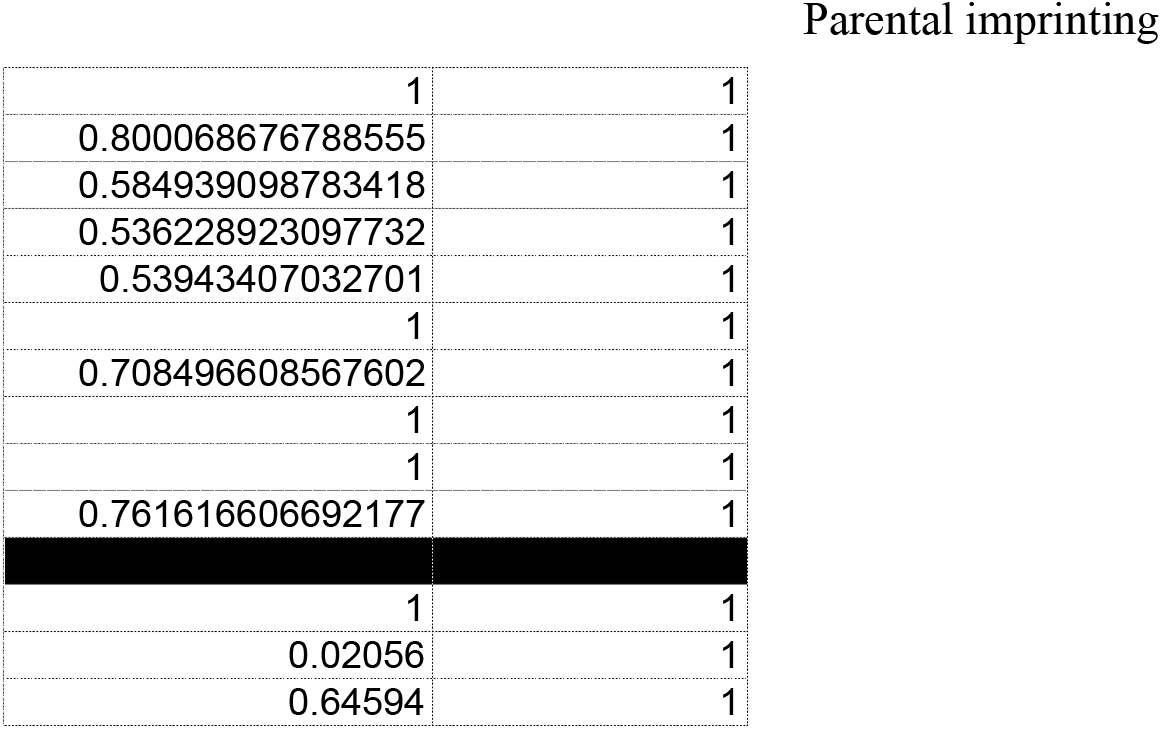

Parental imprinting
| Comparison between tripli or more copies and dupli or less copies |  |
| --- | --- |
| Hypergeometric test p-value | Hypergeometric test FDR (Benjamini-Hochberg procedure) |
| 0.919697883721407 | 1 |
| 0.591671556911165 | 1 |
| 0.731618535243889 | 1 |
| 0.655280844788823 | 1 |
| 0.55040331717341 | 1 |
| 0.89300832429622 | 1 |
| 0.524685862083034 | 1 |
| 0.714964489765281 | 1 |
| 0.82750852910998 | 1 |
| 1 | 1 |
| 0.324438851863797 | 1 |
| 1 | 1 |
| 1 | 1 |
| 0.832830330656687 | 1 |
| 0.296585625240612 | 1 |
| 1 | 1 |
| 0.482545496528408 | 1 |
| 0.173894245758489 | 1 |
| 0.575679270258282 | 1 |
| 0.807846579527661 | 1 |
| 0.831273389308077 | 1 |
| 0.240387740607254 | 1 |
| 0.0626763623704843 | 1 |
| 0.303860354974478 | 1 |
| 0.103473462396853 | 1 |
| 0.020560745644604 | 1 |
| 0.171765715980766 | 1 |
| 0.0624317485962584 | 1 |
| 0.329504317689732 | 1 |
| 1 | 1 |
| 0.389906623547002 | 1 |
| 0.636665677356488 | 1 |
| 0.544224894048767 | 1 |
| 0.325558848240955 | 1 |
| 0.583885952588294 | 1 |
| 0.853047315048397 | 1 |
| 1 | 1 |
| 0.780655009805815 | 1 |
| 0.511810362172052 | 1 |
| 0.466686896846573 | 1 |
| 0.567497718467925 | 1 |
| 0.69752313275528 | 1 |
| 0.862731684825785 | 1 |
| 0.930658132592561 | 1 |
| 1 | 1 |
| 0.937562813354077 | 1 |
| 0.770725733121052 | 1 |
| Parental imprinting |  |
| 1 | 1 |
| 0.800068676788555 | 1 |
| 0.584939098783418 | 1 |
| 0.536228923097732 | 1 |
| 0.53943407032701 | 1 |
| 1 | 1 |
| 0.708496608567602 | 1 |
| 1 | 1 |
| 1 | 1 |
| 0.761616606692177 | 1 |
| 1 | 1 |
| 0.02056 | 1 |
| 0.64594 | 1 |

